# Delayed auditory encoding and variable representation of stimulus regularity in cerebellar lesion patients

**DOI:** 10.1101/2021.02.06.430035

**Authors:** Michael Schwartze, Sonja A. Kotz

**Author notes:** Corresponding author: Faculty of Psychology and Neuroscience, Dept. of Neuropsychology and Psychopharmacology, P.O. Box 616, 6200 MD Maastricht, Netherlands.

## Abstract

The dynamic and fleeting nature of sound necessitates the rapid encoding and use of information distributed over time. Here we investigated cerebellar contributions to these abilities. We measured EEG from cerebellar patients and healthy controls while they listened to “oddball” sound sequences consisting of infrequent pitch-deviant and frequent standard tones. Inter-stimulus-intervals were temporally regular (600 ms) or irregular (200-1000 ms). This allowed probing early event-related potentials (ERP; P50, N100) that reflect repetitive and changing stimulus characteristics in temporally regular or less (irregular) predictable sequences. Further, time-frequency data provided an index of temporal processing variability at the stimulation frequencies. We expected that cerebellar lesions lead to aberrant encoding and use of auditory information, reflected in the ERP morphology of peak amplitudes, latencies and typical suppression effects linked to stimulus predictability. Results confirm longer P50 peak latencies in patients and variable processing at stimulation frequencies covarying with the location of cerebellar damage. These findings further support the idea that the cerebellum might play a generalizable role in the encoding of auditory stimulation over time.

## 1. Introduction

Key human functions such as speech and music build on complex sound patterns unfolding in time and across different timescales. Listening requires rapid and precise encoding of the type and timing of these patterns before the transient signal vanishes. However, the human brain does not just passively react to the sensory environment but modulates its functioning and shapes how input is perceived based on predictions about future sensory events to continuously adapt to a dynamic environment (Friston, 2005; Bar, 2007; Raichle, 2010).

One influential framework that includes a neurofunctionally specific account of how the brain may achieve predictive adaptation is the cerebellar forward model (Wolpert and Miall, 1996; Wolpert et al., 1998). Originally, this framework proposed that the cerebellum might contain multiple forward models of the motor system to facilitate motor control (Wolpert et al., 1998). The forward model is essentially a type of internal model that specifies correct input-output mappings, i.e., predictions about the ideal sensory consequences of movement, and a form of error signal that is used for learning (Ramnani, 2006). This original motor conception was further extended to cognitive functions (Ramnani, 2006; Ito, 2006), thereby specifying potential cerebellar contributions to non-motor behavior (Stoodly and Schmahmann, 2009; Strick et al., 2009; Baumann et al., 2015). It is long known that the cerebellum plays a role in the learning and preparing of sensory events (Bower, 1995; Gao et al., 1996) and the purely sensory processing of auditory information (Petacchi et al., 2005). Similarly, cerebellar contributions to temporal processing in motor production and perception are long recognized (Ivry and Keele, 1989). The idea is that the cerebellum establishes a precise representation of the temporal structure of events in the hundreds-of-milliseconds range (Ivry and Schlerf, 2008; Spencer and Ivry, 2013). The basic concept of the cerebellum as a temporal processing system has recently been refined and extended by integrating functional cerebellar regions with thalamo-cortico-striatal circuits as well as supplementary motor and prefrontal cortices (Schwartze and Kotz, 2013; Petter et al., 2016; Bareš et al., 2019). Such interfacing circuitry may be subserved by structural connections between non-motor regions of the dentate nucleus, the main output relay of the cerebellum with these structures (Bostan and Strick, 2010). Considering the critical role of time and temporal event structure in audition, proposals on temporal processing and predictive adaptation suggest cerebellar contributions to the processing of auditory events, which may specify the temporal regularity of sound events (temporal predictability). Consequently, the cerebellum may play an important role in timing and maintaining predictive activity to guide perception and the allocation of cognitive resources such as attention (Ghajar and Ivry, 2009).

Electroencephalography (EEG) offers a means to investigate different stages of auditory processing with excellent temporal resolution. Most often used as a marker of change detection, are well-established event-related potential (ERP) components such as the mismatch negativity (MMN, elicited by any discriminable change in auditory stimulation and some higher-order cognitive processes (Näätänen et al., 2007)) or the P3 (early on interpreted as an index of surprise and the updating of a mental model of the environment (Donchin, 1981; Linden, 2005)). Here, the processing of change can be considered as a perceptual process, probing stimulus encoding, (sensory) memory, or the allocation of attention.

EEG studies with patients affected by cerebellar degeneration or lesions have shown longer MMN latencies for duration deviants (Moberget et al., 2008) and affected model updating in response to pitch deviants as indexed by the P3b (Kotz et al., 2014). These results indicate that cerebellar damage impacts deviance processing as a basic aspect of auditory cognition. Similarly, application of tDCS over the cerebellum led to a reduced auditory P3b amplitude (Rufener et al., 2020). However, earlier ERP components (P1, N1, P2) and MMN responses to other forms of deviation, including pitch, were found to be comparable to controls (Moberget et al., 2008), establishing an overall heterogeneous picture.

Thus, we explored how diffuse cerebellar lesions alter two ERP components, the P50 and N100, to define how the cerebellum contributes to the temporal expectation of pitch characteristics in early auditory processing (Lange, 2009; Bendixen et al., 2012). We expected to observe modulated P50 and N100 morphology due to cerebellar lesions. To this end, cerebellar patients and healthy controls listened to two continuous tone sequences composed of frequent standard and randomly interspersed infrequent deviant pure tones. Deviants differed from standards in pitch. The two tone sequences differed in stimulus timing, which was fixed in one sequence and randomized in the other, effectively generating one temporally predictable and one less predictable stimulus sequence. This allowed the parallel investigation of encoding of stimulus type (formal structure) and stimulus timing (temporal structure). With this previously established setting (Schwartze et al., 2013; Kotz et al., 2014; Rufener et al., 2020), we expected to replicate earlier findings, i.e., ERP amplitude suppression that reflects contrasts in formal structure and temporal structure. As prior results did not reveal an interaction of formal and temporal structure in early auditory processing (P50, N100; Schwartze et al., 2013), we did not expect such an interaction here. However, we expected that cerebellar damage would affect the morphology of P50 and N100 in terms of (peak) amplitudes and latencies. We hypothesized that cerebellar damage would result in diminished amplitude suppression effects and potentially in longer ERP peak latencies. Such pattern would substantiate the proposed cerebellar role in sensory processing in concert with cortical areas. The specific composition of the stimulus sequences furthermore allowed assessing the proposed role of the cerebellum in the precise encoding of temporal structure on the basis of continuous oscillatory activity at stimulation frequencies in the regular condition. Here, cerebellar damage was expected to increase variability of activity at these frequencies as an indication of imprecise temporal processing.

## 2. Methods

The current study investigated the effects of cerebellar damage on the P50 and N100 ERP components and in the time-frequency domain (power variability at stimulation frequencies), obtained by means of an established experimental paradigm (Schwartze et al., 2013; for additional detail regarding the cerebellar patient population and an analysis of longer-latency ERP components see Kotz et al., 2014).

### 2.1 Participants

A group of eleven cerebellar patients (Fig. 1, Tab. 1) and eleven healthy controls (matching the patients for age, gender, handedness, and education) participated. All participants were informed about the study upon invitation, and provided their informed written consent prior to the recordings. The study was approved by the ethics committee of the University of Leipzig.

**Tab 1.**
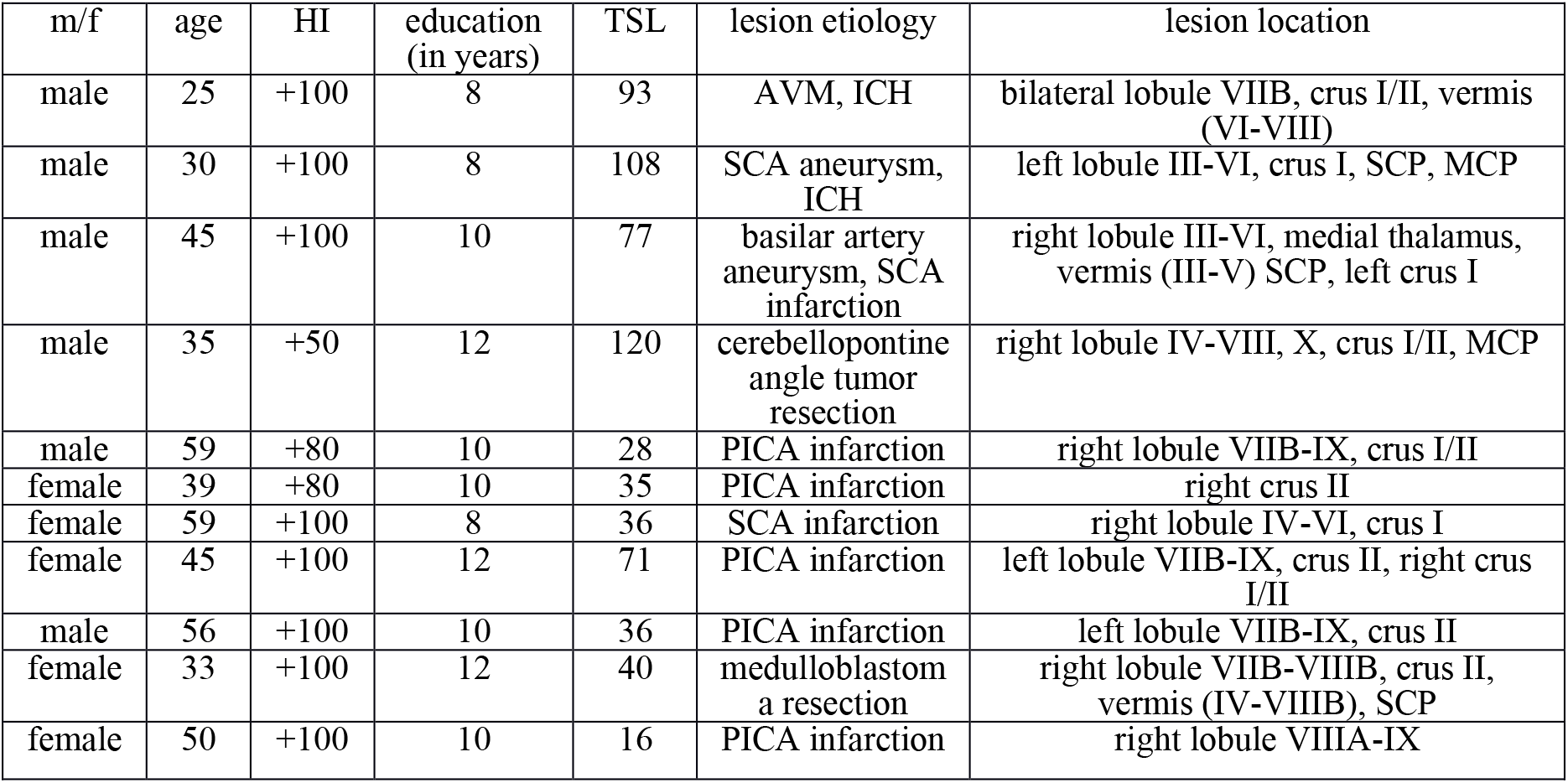
Patient characteristics (adapted from Kotz et al., 2014). A group of eleven cerebellar patients participated in the study. HI = handedness index according to Edinburgh Handedness Inventory (Oldfield, 1971), TSL = time since lesion (in months at the time of testing). Lesion aetiology: AVM = arteriovenous malformation, SCA = superior cerebellar artery, PICA = posterior inferior cerebellar artery, ICH = intracranial haemorrhage. Descriptions of lesion locations refer to the MRI atlas nomenclature of the human cerebellum (Schmahmann et al., 1999), MCP = middle cerebellar peduncle, SCP = superior cerebellar peduncle.

**Fig 1.**
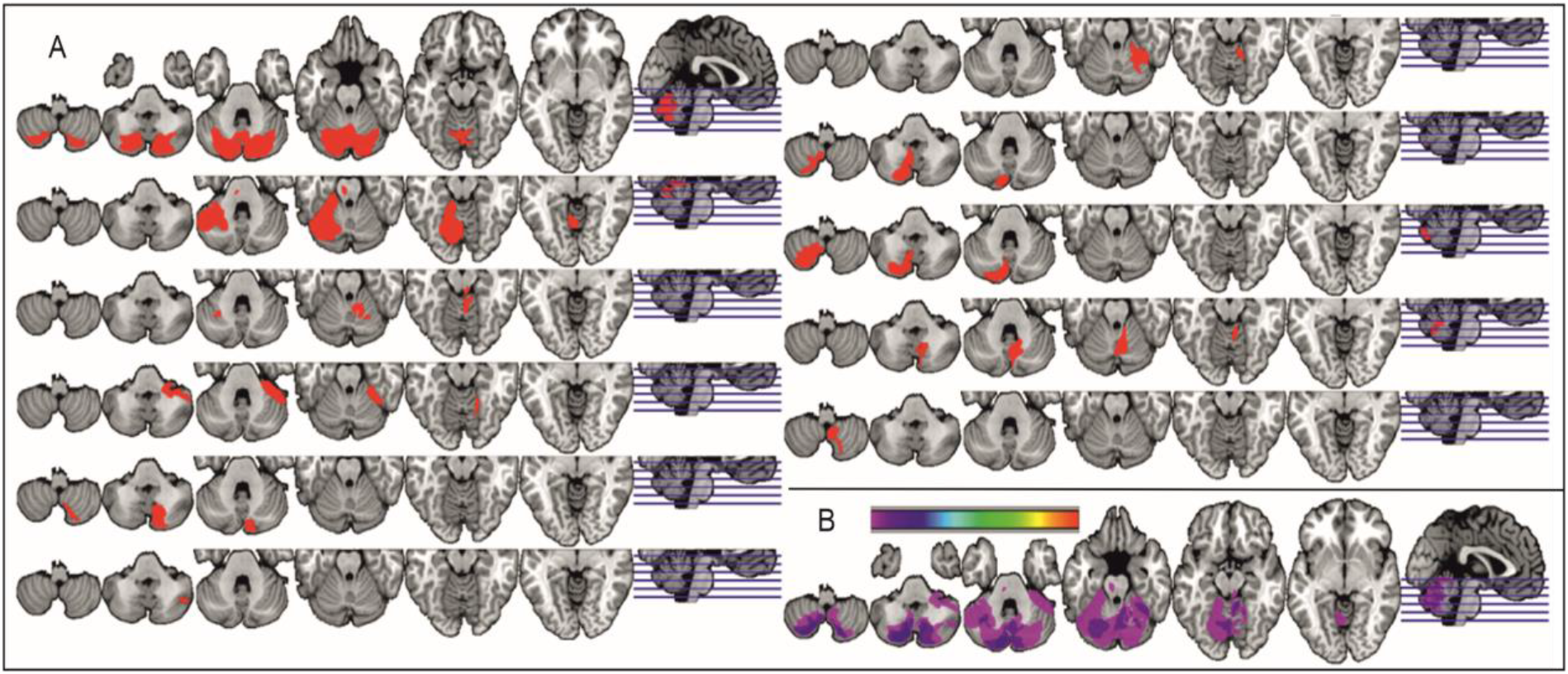
Cerebellar lesions. All lesion locations were hand-drawn into individual anatomical MRI scans using the MRIcron software package (Rorden and Brett, 2000). (A) Individual lesion locations (red) of all patients were then plotted onto an MNI compatible anatomical underlay (Colin27_T1_seg_MNI.nii). (B) An overlay image of the lesion locations was generated, with purple shades reflecting minimal lesion overlap and blue shades maximal lesion overlap (maximum N = 3).

### 2.2 Experimental setup

Participants listened to two continuous sequences of equidurational (300 ms, 10 ms linear rise and fall times) sinusoidal tones (Fig. 2). The presentation order alternated within both participant groups. The sequences were composed of one standard (N=360, 600 Hz) and one deviant (N=90, 660 Hz) tone, corresponding to a standard-to-deviant ratio of 4:1. At the beginning of each sequence, four standard tones were presented to establish a sensory memory trace. Deviant tones were pseudo-randomly presented among standards with no more than two deviants presented in a row. This was achieved by virtually dividing the sequence into groups of four tones, either consisting of standard tones or of three standard and one deviant tone. The position of a deviant tone was counterbalanced across the four possible positions. These groups served as epochs for the time-frequency analyses. However, their presentation was fully randomized, giving participants the impression of listening to one continuous sequence. One of the two sequences was regular (temporally predictable) as the stimulus-onset-asynchrony (SOA) was fixed (900 ms) with an inter-stimulus-interval of 600 ms. The other sequence was irregular (temporally unpredictable; with the restriction that a tone would be presented after an SOA of 1300 ms), with inter-stimulus-intervals randomly chosen from a range of 200-1000 ms.

**Fig 2.**
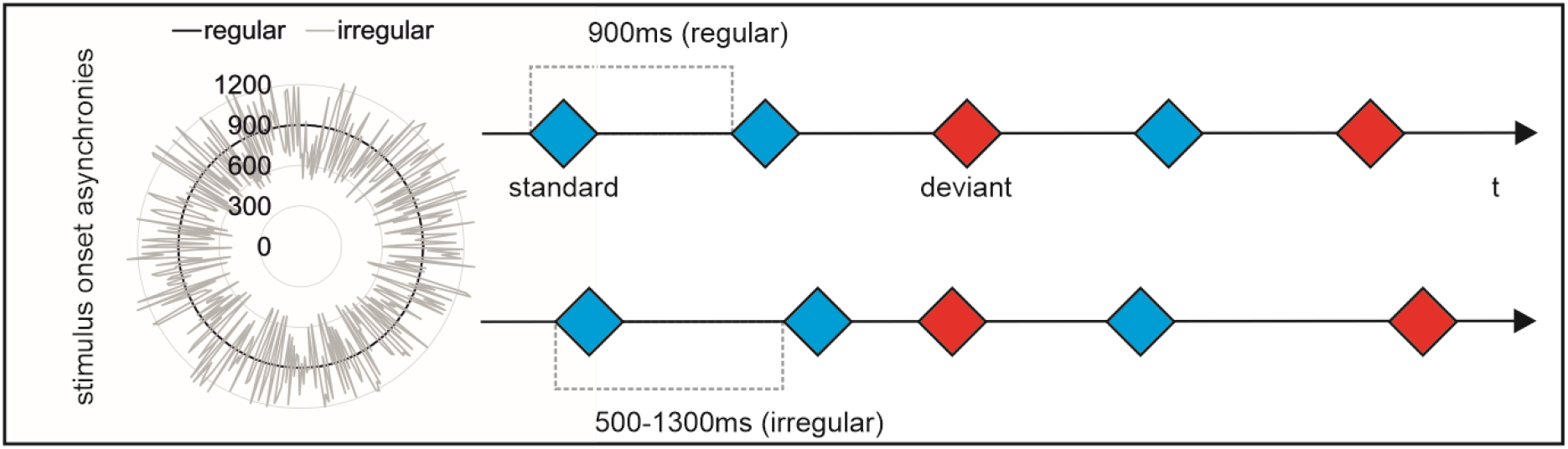
Experimental conditions. Continuous EEG was recorded while participants listened to sequences of 450 equidurational (300 ms) sinusoidal standard (N=360, 600 Hz, blue) and deviant (N=90, 660 Hz, red) tones, presented in two conditions. In the regular condition, tones were separated by fixed inter-stimulus-intervals of 600 ms (900 ms stimulus-onset-asynchronies). In the irregular condition, the same tones were separated by intervals that were randomly chosen from a range between 200 and 1000 ms for each participant (single participant example on the left).

### 2.3 EEG recordings and analyses

Continuous EEG and additional horizontal and vertical electrooculography (EOG) data were recorded from 25 scalp electrodes mounted in an elastic cap and four additional electrodes placed next to the outer canthus of both eyes and above and below the right eye using a sampling rate of 500 Hz, a mastoid reference, and ground placed on the sternum. During the recordings, participants sat comfortably in a dimly lit sound-attenuated room and looked at a computer screen in front of them that displayed an asterisk throughout the recording session. In order to direct attention to the stimulation and to avoid overt motor behavior, thus avoiding additional load on cerebellar patients, participants were asked to silently count the deviant tones embedded in each sequence. For the current analysis of early ERP components and oscillatory activity, data processing was performed with dedicated pipelines using the letswave6 toolbox (https://www.letswave.org) running in Matlab (Mathworks). For ERP analyses, raw data were re-referenced to linked mastoids and bandpass-filtered (Butterworth, 2.0 -30 Hz, order 4). The setting for the high-pass filter was considered adequate for the planned analyses restricted to early ERP components. Independent component analysis (ICA, Runica algorithm) was applied to identify and subsequently remove parts of the data dominated by eye blink- and movement-related artifacts. Data were segmented into epochs lasting from -75 ms to 175 ms relative to tone onset, and a baseline correction was applied. Automatic epoch rejection was performed using an amplitude criterion of 60 µV across all electrodes. Some additional epochs, primarily containing slow drifts (≤ 3 for deviants, ≤ 12 for standards across all participants), were rejected after visual data inspection. The remaining epochs were first averaged per condition for each participant and then for both groups (Fig. 2). This procedure kept a minimum of 80 (out of 90) epochs for deviants and 318 (out of 360) for standards across controls (regular deviants mean: 85.2, SD: 0.4; regular standards mean: 343.4, SD: 2.3; irregular deviants mean: 85.0, SD: 0.2; irregular standards mean: 348.1, SD: 1.6) and patients (regular deviants mean: 84.2, SD: 1.2; regular standards mean: 344.4, SD: 6.0; irregular deviants mean: 84.5, SD: 1.3; irregular standards mean: 342.7, SD: 8.4). For the time-frequency analyses, raw data were bandpass-filtered (Butterworth, 0.3 -30 Hz, order 4). ICA was used to remove data dominated by artifacts. Data were segmented into epochs corresponding to the presentation of a group of four stimuli and the respective inter-stimulus-intervals (3.6 s). Linear detrending was applied for the whole epoch and a 40 µV automatic rejection criterion was applied to an averaged fronto-central region of interest (ROI) channel (see statistical analyses). Finally, the fast continuous wavelet transform (CWT) as provided by letswave6 was performed for a range from 0.5 to 8 Hz (450 lines). This range covered three frequency bands of interest, corresponding to the stimulus-onset asynchrony (SOA, 900 ms, 1.1 Hz), inter-stimulus-interval (ISI, 600 ms, 2.2. Hz), and the stimulus duration (300 ms, 3.3 Hz) employed in the regular condition. Oscillatory activity and its variability at these frequencies was considered indicative of the quality of temporal processing at stimulation frequencies that correspond to the temporal structure of the regular condition.

Statistical analyses were performed by means of SPSS 25 (IBM) on three variables of interest, including peak ERP amplitudes, the corresponding ERP peak latencies, and the variability (SD) of oscillatory power at stimulation frequencies. The variables were derived from one ROI and time-windows lasting from 40 to 80 ms (P50), from 90 to 140 ms (N100), and the whole epoch for time-frequency data. The ROI covered eight fronto-central electrode positions (F3, FZ, F4, FC3, FC4, C3, CZ, C4). These electrodes were selected on the basis of the typical fronto-central scalp topography of the P50 and N100 components. ERP data were analysed by means of 2 x 2 x 2 ANOVAs for both variables, comprising the between-subjects factor *group* (patients vs. controls) and the within-subjects factors *temporal structure* (regular vs. irregular sequence timing) and *formal structure* (standard vs. deviant tone). Time-frequency data were analyzed using a 2 x 2 x 3 ANOVA with the factors *group, temporal structure* and the within-subjects factor *frequency band* (SOA vs. ISI vs. Stimulus). Analyses were performed on z-transformed mean and SD values, focusing on potential differences between the groups, i.e., main effects and interactions involving the factor *group* in order to assess the effects of cerebellar damage on auditory processing. Additionally, the total lesion volume (in cm^3^) and the corresponding centre of mass (in terms of stereotactic coordinates derived from MRIcron, providing an estimation of location) were selected for correlation analyses planned for ERP and time-frequency results that would indicate significant group differences.

## 3. Results

Visual inspection of the ERP data suggested that the expected amplitude suppression pattern was present for the P50 and N100, i.e., there were larger amplitudes in response to deviants than standards and for temporally irregular than regular stimulation (Fig. 3). This pattern was observed for patients and for controls. ERP amplitudes appeared overall smaller in the patient group though.

**Fig 3.**
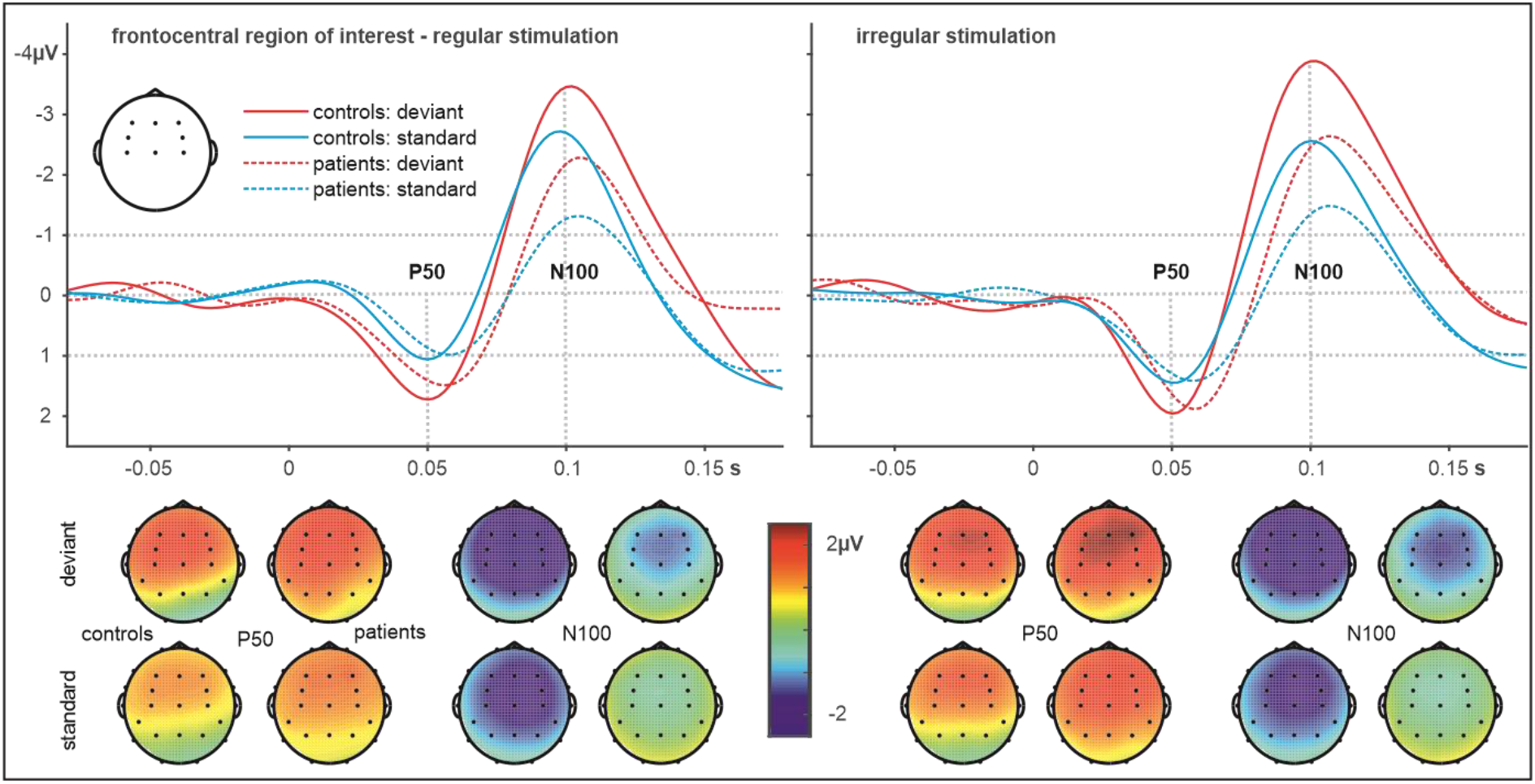
Event-related potential data. P50 and N1 components and corresponding topographical scalp distributions for controls and cerebellar patients as obtained in response to standard and deviant stimuli presented with either regular or irregular temporal structure. A 20 Hz low-pass filter was applied to the data for graphical display only.

The initial visual impression was confirmed by the ANOVA, which yielded significant independent main effects of *temporal structure F*(1.20) = 4.73, *p* = .042, *ηp*^*2*^ = .19 and *formal structure F*(1.20) = 23.60, *p* = .00, *ηp*^*2*^ = .541 for P50 (all other *p* > .26) and of *formal structure F*(1.20) = 42.68, *p* = .00, *ηp*^*2*^ = .68 for N100 peak amplitudes. Notably, the effect of *temporal structure* did not reach significance for the N100 *F*(1.20) = 2.72, *p* = .12, *ηp*^*2*^ = .12). There was a weak trend for an effect of *group F*(1.20) = 2.88, *p* = .11, *ηp*^*2*^ = .13 and for an interaction of *temporal structure* and *formal structure F*(1.20) = 2.34, *p* = .14, *ηp*^*2*^ = .11) that remained non-significant (all other *p* > .48). Overall, these results align with earlier findings but do not confirm an impact of cerebellar damage on P50 and N100 amplitudes (Schwartze et al., 2013). However, the corresponding analysis of peak latencies derived from the P50 time-window yielded a significant main effect of *group F*(1.20) = 8.73, *p* = .01, *ηp*^*2*^ = .30 (all other *p* > .18), indicating globally and consistently longer latencies in patients for regular deviants (Mean 57 (SD 11) ms; controls 52 (8) ms) and standards (60 (14) ms, controls 48 (8) ms) as well as for irregular deviants (58 (9), controls 48 (8) ms) and standards (58 (15) ms, controls 53 (12) ms). Although the same pattern of delays emerged, this difference was not significant for N100 latencies that rather revealed a trend interaction of temporal and formal structure *F*(1.20) = 2.25, *p* = .15, *ηp*^*2*^ = .26; overall there were no significant findings in this case (all other *p* > .26) though.

The ANOVAs (Greenhouse-Geisser correction where applicable) on the z-transformed mean and SD of the time-frequency data (Fig. 4) yielded no significant findings for means (all *p* > .16) but a significant interaction *group* x *temporal structure F*(1.20) = 7.87, *p* = .01, *ηp*^*2*^ = .28 next to a non-significant trend for an interaction of all three factors for SD values *F*(2.40) = 3.08, *p* = .07, *ηp*^*2*^ = .13. Step-down analyses of the *group* x *temporal structure* interaction by the factor *temporal structure* revealed a significant group difference for the regular *F*(1.20) = 4.82, *p* = .04, *ηp*^*2*^ = .19 but not for the irregular condition *F*(1.20) = .31, *p* = .59, *ηp*^*2*^ = .02.

**Fig 4.**
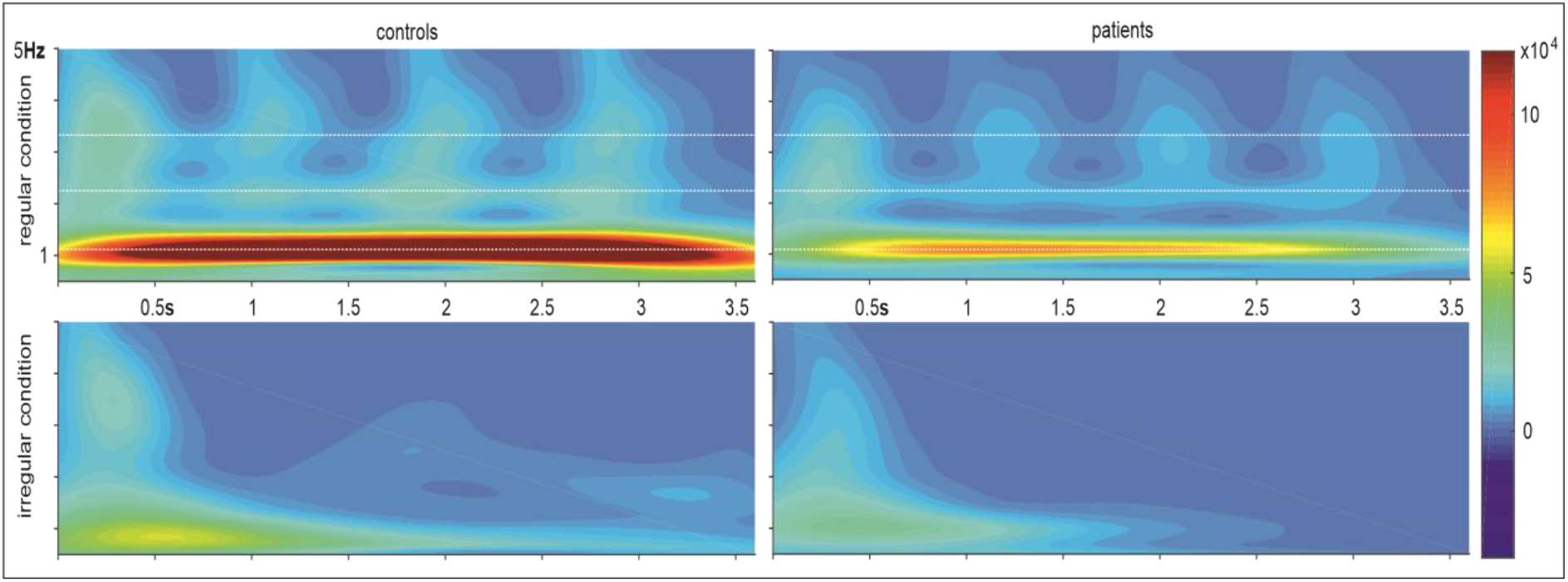
Time-frequency data. Variability (SD) of oscillatory activity (power) was taken as an indication of temporal processing consistency and derived for three frequencies of interest (white lines), reflecting the temporal structure of the continuous regular stimulation (stimulus-onset asynchrony (900 ms, 1.1 Hz), inter-stimulus interval (600 ms, 2.2 Hz) and stimulus duration (300 ms, 3.3 Hz)).

To test for potential effects of lesion extent and location on the ERP latency group difference, the average of the respective peak latencies across the P50 ERPs was correlated with the lesion volume and stereotactic (x/y/z) coordinates of the centre of mass for each lesion. This procedure yielded no significant result. However, the same approach provided a significant positive correlation (Fig. 5) between the average time-frequency variability across frequencies in the regular condition and x-coordinates (lateral dimension) (*r* = .61, *p* = .047), indicating that further right-lateralized lesions were associated with increased variability, thereby establishing a direct link between electrophysiological data and type of cerebellar damage.

**Fig 5.**
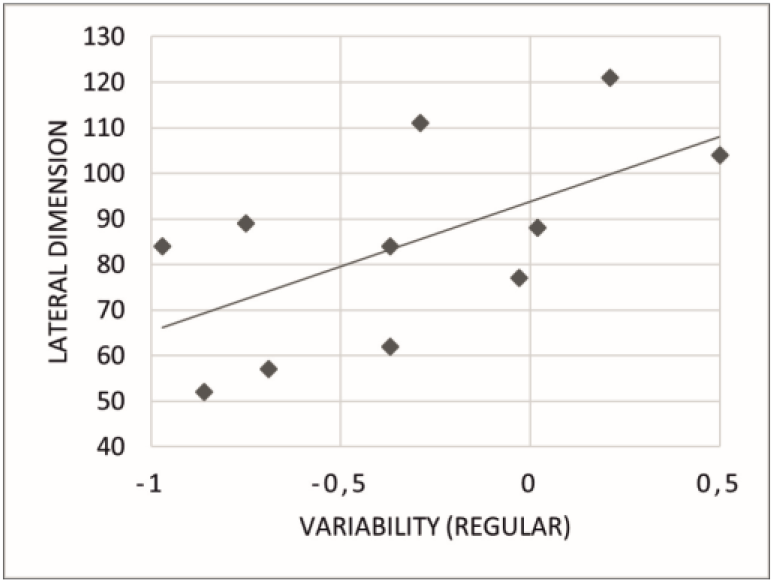
Correlation result. Average variability across frequencies in the regular condition (z-transformed SD) was positively correlated with the lateral dimension (x-coordinates) of cerebellar damage, indicating that damage located further to the right was associated with increased variability.

## 4. Discussion

The predictive coding account to brain function has become a prevalent approach in the study of auditory perception (Heilbron and Chait, 2018; Denham and Winkler, 2020). However, to generate temporal predictions, perception has to rely on some form of an adequately precise representation of the temporal structure of the dynamic input. This may be achieved in interaction with temporal processing mechanisms that are associated with sensorimotor brain structures, including the cerebellum (Merchant et al., 2013; Teghil et al., 2019). The current study investigated the effect of cerebellar damage on electroencephalographic markers of auditory perception, namely on the P50 and N100 ERP components and on the variability of oscillatory activity at stimulation frequencies. Cerebellar damage was found to affect both measures.

The ERP results replicated earlier findings in healthy young participants confirming P50 and N100 suppression effects (Schwartze et al., 2013). In line with other reports (e.g., Lange, 2009), P50 and N100 amplitudes were smaller for regular than irregular stimuli. This was the case for predictability of formal structure (frequent standard vs. infrequent deviant tones) and temporal structure (regular vs. irregular stimulus timing). However, the effect of temporal structure did ultimately not lead to N100 amplitude suppression statistically. Also, in line with earlier patient findings (Moberget et al., 2008), there were no qualified ERP amplitude differences between the patients and controls. Patients in the current study showed longer ERP peak latencies. While attempting to link these latency differences to lesion volume and centres of mass, this analysis failed to produce significant results. However, an overall group difference suggests a generalizable contribution of the cerebellum to the encoding of auditory stimuli over time. The prolonged ERP latencies may be indicative of a transmission delay of signals to cortical targets involved in the generation of the EEG signal as shown in the fronto-central ROI. Rapid signal transmission from the cerebellum may serve to initiate temporal processing in cortical targets that include the supplementary motor area (Schwartze and Kotz, 2013; Patter et al., 2016). Next to temporo-parietal generators, there is evidence that the P50 also has frontal sources that may include the supplementary motor area (Grunwald et al., 2003; Korzyukov et al., 2007; Kurthen et al., 2007). Similar findings have previously been discussed for the N100 (Alcani et al., 1994; Giard et al., 1994). If the cerebellar temporal processing system interfaces with other structures such as supplementary motor and prefrontal cortical areas in audition, the current results may indicate a constant delay in an initiation mechanism, which may also explain why peak amplitudes were not significantly different between groups.

Despite such considerations and other proposals that discuss a role of the cerebellum in temporal processing and auditory function in general (Petacchi et al., 2005; Parsons et al., 2009), it is still largely unclear if and how the cerebellar temporal processing system interfaces with auditory processing, how it may factor into well-established characteristics of auditory processing (e.g., in terms of cortical functional lateralization), and moreover, how event-timing and associated neural circuitry may relate to adaptive and predictive sensorimotor mechanisms as basic as Pavlovian conditioning (Friston and Herreros, 2016). To further differentiate the role of the cerebellum in temporally predictive adaptation, it may be relevant to re-assess earlier comparative structural and functional evidence concerning auditory pathways through the cerebellar dentate and rostral thalamus to motor and frontal cortices that may allow rapid encoding and transmission of signals for conditioning (Woody et al., 1998; Petacchi et al., 2005).

Patients did not differ from controls in their ERP responses to temporal structure, i.e., the difference between temporally fully predictable and less predictable stimulus timing. In contrast, controls and patients differed in terms of variability of oscillatory activity in the regular condition, suggesting a less consistent representation of the temporal structure of the regular stimulation in patients. The additional finding that more right lateralized damage was associated with higher variability is relevant in light of the mainly contralateral connectivity of the cerebellum and association cortex (Wang et al., 2013). Although it is important to note that the found correlation does not imply right-hemisphere lesions, the result is at least compatible with the finding of a left supplementary motor area preference for subsecond timing (Schwartze et al., 2012), potentially indicating contralateral cerebello-cortical interaction.

Considering the proposed role of the cerebellum in event-timing, the negative ERP finding seems surprising, especially as patients with cerebellar degeneration did not show the typical suppression effect associated with periodic stimulation (Moberget et al., 2008). Although these negative findings have to be interpreted with care, the question arises if the role of the cerebellum is primarily related to the precise and potentially automatic encoding of temporal event structure (Spencer and Ivry, 2013). If this is the case, the temporal processing capacities of the basal ganglia and supplementary motor cortices (Merchant et al., 2013; Petter et al., 2016) may be instrumental for the actual recognition and subsequent use of a temporal pattern in cognitive operations. As these structures are intact in the patient population, these capacities may partly compensate for the consequences of cerebellar damage. In line with this proposal, basal ganglia lesion patients tested with the identical paradigm showed a selective indifference towards the manipulation of temporal regularity in their P50 response (Schwartze et al., 2015). However, the current findings concerning ERP latencies in combination with earlier reports (Kotz et al., 2014), confirm an impact of cerebellar damage on auditory processing, and potentially on different stages of auditory processing. While temporal predictability and deviance processing interact at later processing stages as reflected in the P300, and more specifically the P3b component, early processes as reflected in the P50 and N100 seem to encode formal and temporal structure more independently. Although these findings are compatible with the proposed function of the cerebellum in purely perceptual auditory processing and temporal processing, it stands to reason how these stages relate to each other functionally, and how the cerebellum is embedded into a wider network, in which different structures may interact to achieve the overarching goal of efficient predictive adaptation to the environment.

## Acknowledgements

The authors would like to thank Anne-Kathrin Franz and Heike Boethel for support during data acquisition, and Anika Stockert for support in the preparation of the clinical data. Part of this work has been conducted while the first author was a member of staff at the Max Planck Institute for Human Cognitive and Brain Sciences in Leipzig, Germany.

